# The Anorexia Nervosa Genetics Initiative: Overview and Methods

**DOI:** 10.1101/234013

**Authors:** Laura M Thornton, Melissa A Munn-Chernoff, Jessica H Baker, Anders Juréus, Richard Parker, Anjali K Henders, Janne T Larsen, Liselotte Petersen, Hunna J Watson, Zeynep Yilmaz, Katherine M Kirk, Scott Gordon, Virpi M Leppä, Felicity C Martin, David C Whiteman, Catherine M Olsen, Thomas M Werge, Nancy L Pedersen, Walter Kaye, Andrew W Bergen, Katherine A Halmi, Michael Strober, Allan S Kaplan, D. Blake Woodside, James Mitchell, Craig L Johnson, Harry Brandt, Steven Crawford, L. John Horwood, Joseph M Boden, John F Pearson, Laramie E Duncan, Jakob Grove, Manuel Mattheisen, Jennifer Jordan, Martin A Kennedy, Andreas Birgegård, Paul Lichtenstein, Claes Norring, Tracey D Wade, Grant W Montgomery, Nicholas G Martin, Mikael Landén, Preben Bo Mortensen, Patrick F Sullivan, Cynthia M Bulik

## Abstract

**Background:** Genetic factors contribute to anorexia nervosa (AN); and the first genome-wide significant locus has been identified. We describe methods and procedures for the Anorexia Nervosa Genetics Initiative (ANGI), an international collaboration designed to rapidly recruit 13000 individuals with AN as well as ancestrally matched controls. We present sample characteristics and the utility of an online eating disorder diagnostic questionnaire suitable for large-scale genetic and population research.

**Methods:** ANGI recruited from the United States (US), Australia/New Zealand (ANZ), Sweden (SE), and Denmark (DK). Recruitment was via national registers (SE, DK); treatment centers (US, ANZ, SE, DK); and social and traditional media (US, ANZ, SE). All cases had a lifetime AN diagnosis based on DSM-IV or ICD-10 criteria (excluding amenorrhea). Recruited controls had no lifetime history of disordered eating behaviors. To assess the positive and negative predictive validity of the online eating disorder questionnaire (ED100K-v1), 109 women also completed the Structured Clinical Interview for DSM-IV (SCID), Module H.

**Results:** Blood samples and clinical information were collected from 13,364 individuals with lifetime AN and from controls. Online diagnostic phenotyping was effective and efficient; the validity of the questionnaire was acceptable.

**Conclusions:** Our multipronged recruitment approach was highly effective for rapid recruitment and can be used as a model for efforts by other groups. High online presence of individuals with AN rendered the Internet/social media a remarkably effective recruitment tool in some countries. ANGI has substantially augmented Psychiatric Genomics Consortium AN sample collection. ANGI is a registered clinical trial: clinicaltrials.gov NCT01916538; https://clinicaltrials.gov/ct2/show/NCT01916538?cond=Anorexia+Nervosa&draw=1&rank=3.

## Introduction

Genetic factors play a substantial role in liability to anorexia nervosa (AN). Relative risk is ~11 in female first-degree relatives of those who have AN (1), and twin studies estimate heritability at 48%-74% (2–5). The Eating Disorders Working Group of the Psychiatric Genomics Consortium (PGC-ED) reported the first genome-wide significant locus on chromosome 12 in a region previously implicated in type 1 diabetes and autoimmune illnesses (6). The goal of the Anorexia Nervosa Genetics Initiative (ANGI) was to rapidly expand available samples for genome-wide association studies (GWAS) of AN. Given the complex genetic architecture of psychiatric disorders, large sample sizes, perhaps hundreds of thousands, are necessary to identify variants associated with these disabling conditions (7, 8). We provide an overview of recruitment procedures and methods used in ANGI, which collected the largest sample of AN cases and controls in the world, considerably augmenting existing samples in the PGC-ED (9).

A secondary goal of ANGI was to provide efficient phenotyping for future investigations. In the absence of biomarkers for psychiatric disorders, the large sample sizes required for GWAS encourage the development of valid assessments with minimal investment of time and effort. Structured clinical interviews, regarded by some as the preferred method for assessing eating disorders (10), are not economically feasible for such studies. Well-validated and easily accessible self-report assessments provide an alternative and may encourage greater openness about disordered eating behaviors than face-to-face interviews (11, 12). To this end, we report the validity of an online eating disorder questionnaire (ED100K-v1) designed to capture AN cases and controls for inclusion in ANGI. Data generated from ANGI will provide pertinent information about the etiology of AN and contribute to the development of biologically informed therapeutics.

## Methods and Materials

### Collaborative Arrangements

ANGI is an international collaboration sponsored by the Klarman Family Foundation and the National Institute of Mental Health. Four primary hubs for data collection, selected because of experience in collecting large genetic samples and access to individuals with a lifetime history of AN, included the University of North Carolina at Chapel Hill [(UNC), United States (US)]; QIMR Berghofer Medical Research Institute [Brisbane, Australia with assistance from the University of Otago in Christchurch, New Zealand (ANZ)]; Karolinska Institutet [Stockholm, Sweden (SE)]; and Aarhus University [Aarhus, Denmark (DK)]. The organizational structure consists of a lead principal investigator (Bulik); site principal investigators (Bulik, Martin, Landén, Mortensen); deputy director (Thornton); steering committee (chair: Bulik); biological sample committee (chair: Sullivan); analysis group (chair: Sullivan); phenotype group (chair: Thornton); and publication and editorial committee (chair: Bulik). A scientific advisory council provided external oversight of study procedures; monitored progress of sample collection, genotyping, and data analysis; and insured adherence to ethical standards and data sharing procedures. The appropriate ethics or institutional review boards at each location approved study protocols.

### General Study Procedures

In the US, ANZ, and SE, inclusion criteria for cases were based on the Diagnostic and Statistical Manual of Mental Disorders, 4^th^ edition (DSM-IV) AN Criteria A, B, and C. Amenorrhea (Criterion D) was not required since it is not applicable to males and was removed from the DSM-5. In DK, cases were defined as any individual present in the national patient register or who presented in the clinic with an ICD-10 diagnosis of F50.0 (AN) or F50.1 (atypical AN).

Recruitment and study procedures varied across sites (see **Figure 1**) due to local ethical requirements or the manner in which cases were identified and are discussed below.

**Figure 1.**
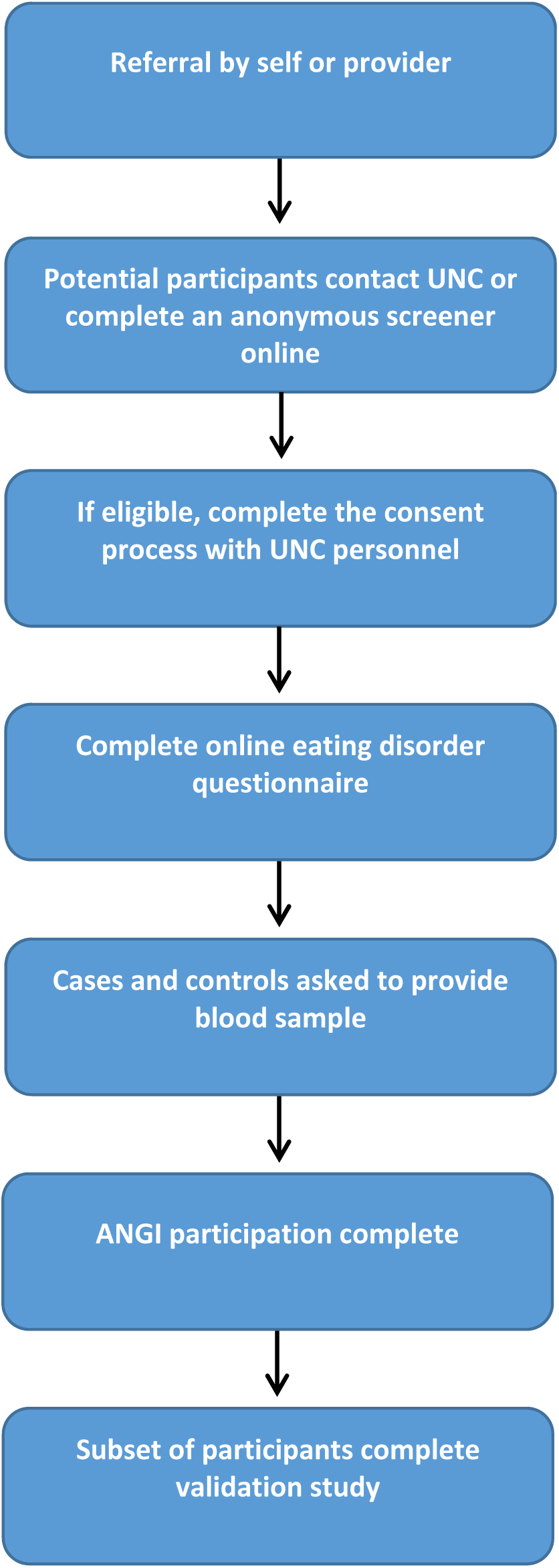
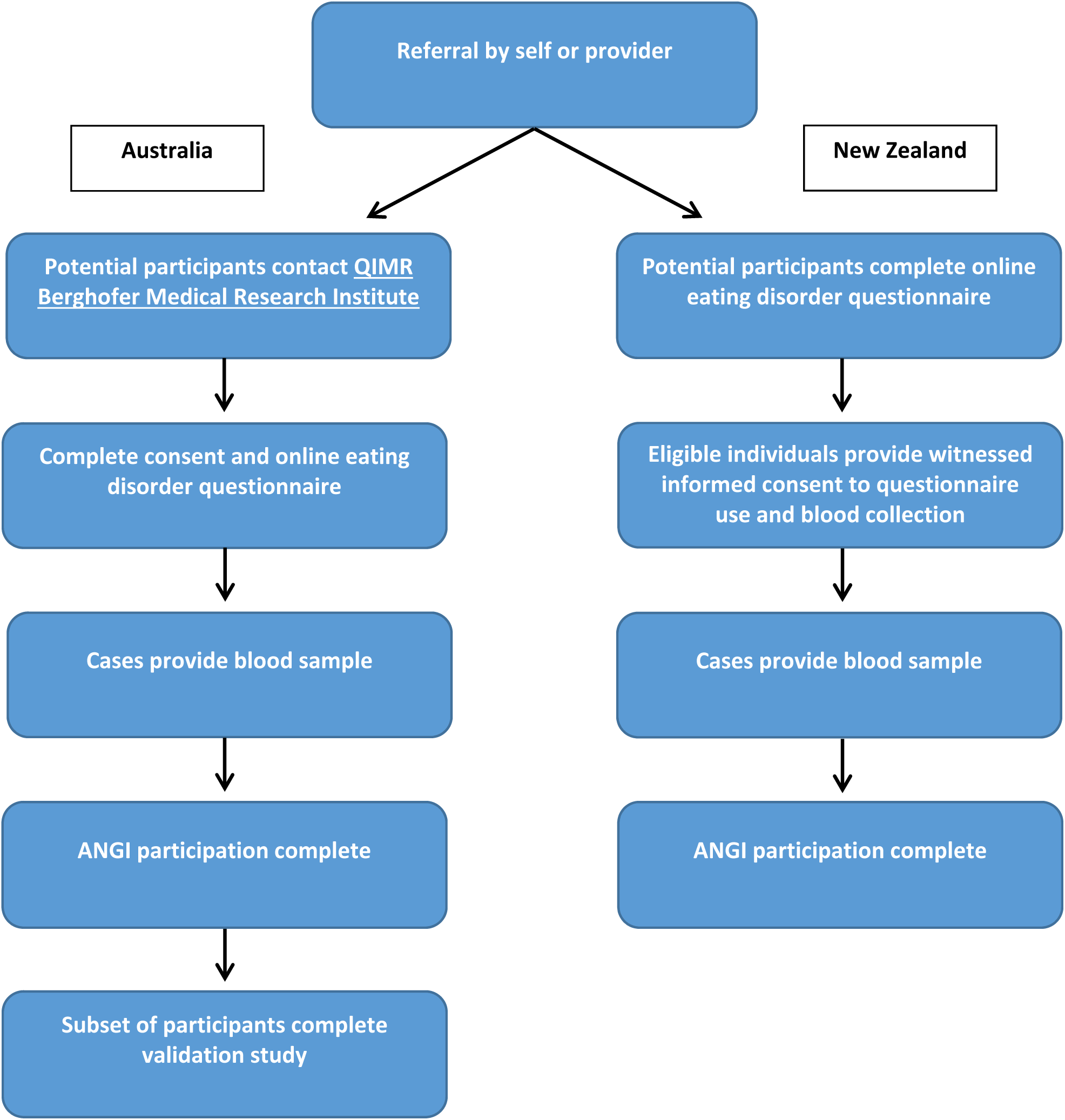
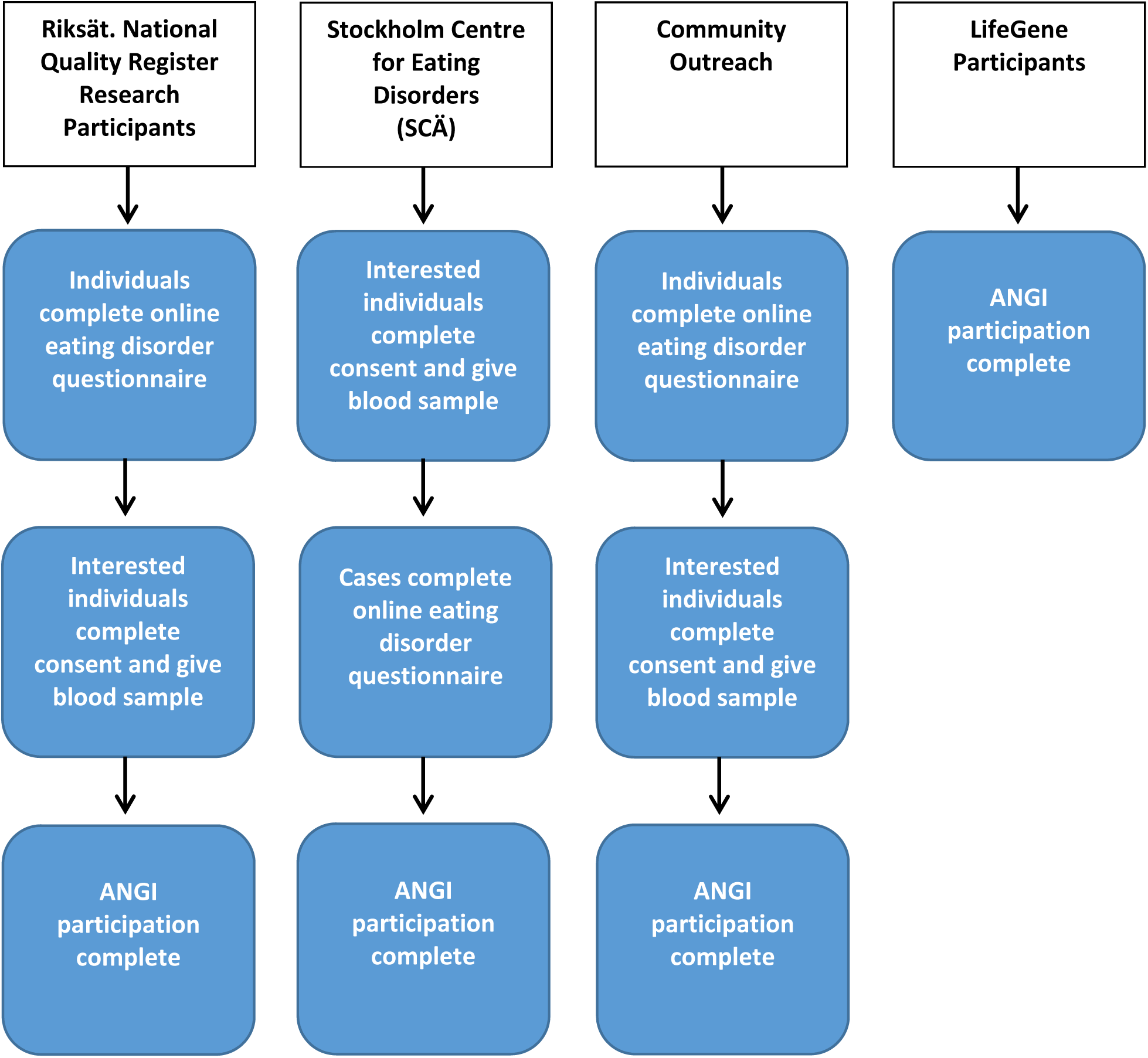
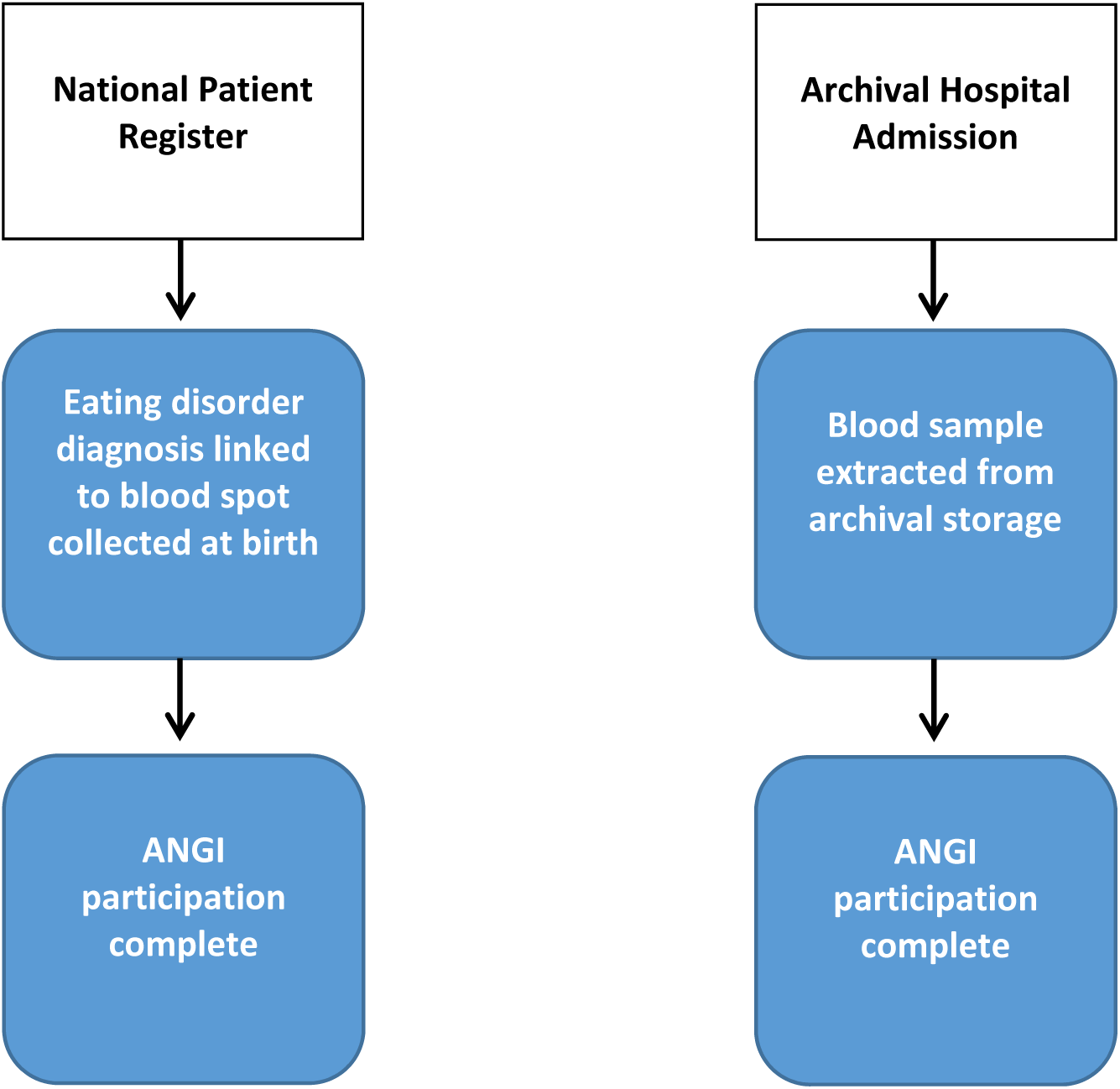
Case recruitment procedures and study design for the Anorexia Nervosa Genetics Initiative (ANGI) by site. a)United States, Case recruitment procedures and study design for the Anorexia Nervosa Genetics Initiative (ANGI) by site. a) Australia / New Zealand, Case recruitment procedures and study design for the Anorexia Nervosa Genetics Initiative (ANGI) by site. c) Sweden, Case recruitment procedures and study design for the Anorexia Nervosa Genetics Initiative (ANGI) by site. b) Denmark

#### United States and Australia/New Zealand

##### Recruitment Approach

A primary focus of our recruitment strategy was to include individuals who: 1) may not live close to recruitment centers, but who desired to participate; 2) may have suffered from AN and never received treatment; 3) have become detached from providers; or 4) are currently in treatment programs. Therefore, we used traditional and novel approaches in the US (ANGI-US) and ANZ [ANGI-ANZ(AUS) and ANGI-ANZ(NZ)]. Recruitment avenues included social and electronic media; eating disorders treatment centers; posts on Recovery Record (a mobile app for individuals in recovery from their eating disorder); prominent eating disorder blogs; and traditional media (e.g., newspaper advertisements and flyers). In the US, ANZ, and SE, television and radio interviews were conducted and website interviews were posted online, which were followed-up on social media sites (e.g., Facebook, Twitter) with links to additional stories. The highly successful Australian media campaign was awarded a commendation in the Public Relations Institute of Australia Queensland Awards for Excellence.

Collecting recruitment information regarding how participants heard about the study (US, ANZ) allowed us to evaluate the relative effectiveness of recruitment strategies for cost-effective implementation in future genetic and non-genetic research. Furthermore, by recruiting from a wide variety of sources, our sample reflects a broad range of individuals who have had AN and not just those seeking treatment. This is critically important as only 33% of individuals with AN receive treatment for the illness (13, 14).

##### Study Procedures

Individuals who resided in the US (ages ≥12 years), AUS (ages ≥13 years), or NZ (ages ≥14) either self-identified or were referred to the study. In the US, individuals first completed an online screener to determine study eligibility. In all three countries, those who were interested in study participation (and deemed eligible by the brief screen in the US) completed the consent process and online questionnaire based on the Structured Clinical Interview for the DSM-IV (SCID) (15) Module H (eating disorders). Core diagnostic questions from the SCID-Module H were adapted for self-report for ANGI (ED100K-v1) to capture information on lifetime history of eating disorders. Once the questionnaire was completed, the participant provided a blood sample (see DNA Sampling).

Controls assessed for ANGI-US were included in the study if they had no lifetime history of any disordered eating behavior and did not meet any criteria for cases. Additional control samples were obtained from The Price Foundation Study AN Trios Study (16, 17) [ANGI-US(PFCG)]. Female controls, ages 18-65 years at assessment, were matched to ANGI-US cases by ancestry. Control women were required to have lifetime adult minimum body mass index (BMI; weight [kilograms] / height^2^ [meters]) > 19 and maximum BMI < 27. They had no history of an eating disorder or disordered eating behaviors, nor any first-degree relative with an eating disorder.

In Australia, data from 17,158 participants from the QSkin Sun and Health Study (18) (QSkin, http://QSkin.qimrberghofer.edu.au/) were made available as controls for ANGI-ANZ(AUS). This study, focusing on skin cancer and its risk factors, is being conducted by the Population Health Department at QIMR (PI: David Whiteman). It uses an electoral roll sample of Queenslanders ages 40-60 years (19). Eating disorders are included in the disease checklist: controls with no history of eating disorders were selected. Consent for QSkin allows for inclusion in other research studies. Additional information about recruitment and study procedures from the ANGI-ANZ(AUS) hub is described elsewhere (20).

Potential controls for participants from New Zealand were made available by the Christchurch Health and Development Study (CHDS, http://www.otago.ac.nz/christchurch/research/healthdevelopment/) (21), a longitudinal study of individuals born in 1977. Controls were selected based on negative responses to a self-report eating disorders screen and availability of genotype information (N=739).

#### Sweden

##### Recruitment Approach

Four recruitment strategies were used for ANGI-SE. 1) Individuals in Riksät-National Quality Register for Eating Disorders Treatment (22) [ANGI-SE(Riksät)], which includes eating disorder information from individuals seeking treatment for an eating disorder in Sweden since 1999, who had been in treatment for AN, bulimia nervosa, or eating disorder not otherwise specified were sent a letter asking them to complete a follow-up questionnaire, which included the ED100K-v1 questionnaire. 2) Study nurses at the Stockholm Centre for Eating Disorders [ANGI-SE(SCÄ)] recruited cases. A research nurse discussed the study and reviewed the consent process with patients with AN who came into the center. When participants consented, a blood sample was taken and the participant was directed to complete the ED100K-v1 questionnaire. 3) We recruited cases and controls from the community using traditional media, social media, and the Swedish ANGI website (www.angi.se), directly linking to the questionnaire [ANGI-SE(Community)]. 4) Cases and controls were identified in LifeGene (23) [ANGI-SE(LifeGene), https://www.lifegene.se/], an ongoing study initiated in 2010 to evaluate how genes, environment, and lifestyle affect health. Individuals who enrolled in LifeGene completed an eating disorder assessment similar to the ED100K-v1 questionnaire and provided a blood sample. Algorithms for case and control identification using the LifeGene eating disorder assessment were harmonized with the ED100K-v1 questionnaire.

##### Study Procedures

From ANGI-SE(Riksät), potential cases (ages 15-80 years), willing to participate in research and identified by meeting ANGI case criteria from the ED100K-v1 questionnaire or with a clinical AN diagnosis, were contacted by a research nurse and invited to participate. If interested, they were sent informed consent papers, a phlebotomy kit, and instructions on how to donate blood at a local laboratory. Cases from ANGI-SE(Community) were contacted by a research nurse and followed the same procedure as Riksät-based recruitment.

Statistics Sweden provided new, population-based controls [ANGI-SE(Community)] who were matched to cases from ANGI-SE(Riksät) on age and sex. Identified individuals were asked about height and weight history, eating disorder history, and history of other psychiatric disorders. Those who had no lifetime history of an eating disorder and agreed to be in the study were sent a blood kit to provide a DNA sample. An additional 3,000 archived control samples were identified from LifeGene [ANGI-SE(LifeGene)]-all screened negative for any eating disorder history.

#### Denmark

##### Recruitment Approach

The primary recruitment approach for ANGI in DK (ANGI-DK) utilized the national register and biobank system. Every person who has had permanent residence or was born in Denmark since April 1, 1968 is assigned a unique person identifier used across all Danish social and health care services. This information can be employed to determine lifetime AN diagnoses based on contacts with the hospital system and can be linked to additional registers. Moreover, phenylketonuria (PKU) cards from birth are stored for all individuals in The Danish Neonatal Screening Biobank (DNSB) at the Staten Serum Institut in Copenhagen and indexed by the personal number (see DNA Sampling). PKU cards for individuals who met inclusion criteria and for controls were extracted from DNSB. Additional samples [ANGI-DK(Clinic)] were obtained from the Danish Psychiatric Biobank that recruits individuals with mental disorders for genetic studies (24). The biobank holds DNA samples and demographic, familial, and clinical information, including complete records of all hospital in-and out-patient admissions and diagnoses.

##### Study Procedures

All cases and controls (birth years 1981-2005) from DK registers were identified from the national patient register. ANGI-DK controls were selected from DNSB (24) based on date of birth, sex, and absence of major psychiatric disorders.

For ANGI-DK(Clinic), cases were defined as patients with at least one recorded hospital admission resulting in the assignment of an ICD-10 diagnosis of F50.0 or F50.1. Clinic cases were women born 1947-1980 (age range: 35-68 years).

### DNA Sampling

For ANGI-US, ANGI-ANZ, and ANGI-SE, each new participant provided blood samples in EDTA tubes. In the US, after questionnaire completion, participants had their blood drawn by a mobile phlebotomy company, UNC, a local laboratory, or a physician’s office. Samples were mailed directly to the National Institute of Mental Health designated laboratory at Rutgers University Cell and DNA Repository (RUCDR), where DNA extraction occurred. Sample collection procedures for ANGI-US(PFCG) are described elsewhere (25).

For ANGI-ANZ(AUS), blood draws occurred in community phlebotomy centers. A prepaid, addressed consignment note for overnight delivery was included. All kits were returned to QIMR for processing. For ANGI-ANZ(NZ), participants across New Zealand first provided witnessed informed consent. They were then sent blood kits which they took to a preferred phlebotomy center where samples were drawn and sent through the national laboratory courier system to Christchurch. Whole blood samples were stored in a tissue bank prior to sending to QIMR to be sent with the ANGI-ANZ(AUS) samples to RUCDR for DNA extraction.

Oragene saliva samples were collected for controls from QSkin. Samples were genotyped at Erasmus University on the Illumina Global Screening Array (GSA) chip. Since all cases had DNA samples extracted from blood and all controls had DNA extracted from saliva, we genotyped DNA from both blood and saliva from 107 cases to compare call rates. Of these,102 samples from blood and 101 samples from saliva (100 overlapping samples) passed genotyping quality tests. The genotypes from blood and saliva are concordant an average of 99.69% of the time for a given person (257,712 tested markers).

Participants in the CHDS provided either peripheral blood (92%) or Oragene saliva (8%) during interviews in 2005. Genotyping of these samples was previously carried out using Illumina 660W Quad chips (26).

For ANGI-SE (Riksät, SCÄ, Community), newly recruited cases and controls provided blood samples at their nearest hospital and mailed them to Karolinska Institutet Biobank. ANGI-SE(LifeGene) samples from cases and controls were also stored at Karolinska Institutet biobank. DNA was extracted at Karolinska Institutet.

For ANGI-US, ANGI-ANZ, and ANGI-SE, all DNA was sent to the Broad Institute for genotyping on the Illumina GSA chip.

For ANGI-DK, DNA from blood spots (which can be linked to other Danish databases discussed above) from birth on all participants were was extracted from two biobanks at the Staten Serum Institut in Copenhagen. The first biobank is the DNSB, which collects blood spots from all children born in Denmark for the purpose of neonatal screening for PKU and other diseases in the newborn (27). The second biobank contains maternal serum samples drawn during week 16 of pregnancy for approximately 20% of the DNSB. Prior research demonstrated that the amplified DNSB sample results are of as high quality as results from conventionally isolated DNA samples for both GWAS genotyping and direct sequencing (28, 29). DNA was sent to the Broad Institute for genotyping on the Illumina PsychArray; 3,769 control samples were genotyped as part of ANGI-DK. Data for the remaining control samples and for 224 cases were supplied by the Lundbeck Initiative for Integrative Psychiatric Research (iPsych) consortium (24) and were genotyped at the same time and on the same platform as those for ANGI-DK. The ANGI-DK(Clinical) DNA samples were extracted from whole blood samples and sent to the Broad for genotyping on the Illumina GSA chip.

### Validity of the Online Eating Disorder Questionnaire (ED100K-v1)

To assess the validity of the ED100K-v1 questionnaire, we completed follow-up phone interviews using the SCID-Module H with a random subsample of ANGI participants who agreed to recontact from ANGI-US and ANGI-ANZ(AUS). The validity sample included 52 women from the US and 58 women from AUS. Data from one individual from the US was removed from analyses because the highest, not the lowest, weight was reported in the SCID-H. Interviews were conducted by a PhD-level clinician or interviewer with experience administering structured and semi-structured psychiatric diagnostic interviews. All interviewers were trained to SCID criteria by a certified PhD-level clinician from the UNC Assessment Core. Training included reviewing SCID protocol and instructions, observing SCID interviews, reviewing video/audio taped interviews to score, and being observed completing a SCID interview by a PhD-level clinician. Interviewers and the Principal Investigator met regularly for consensus.^1^

### Statistical Analyses

Statistical analyses were conducted using SAS version 9.3 (30). We present the percentage of participants who heard about ANGI through each source and temporal data on enrollment relative to media launches. We evaluated the extent to which individuals enrolled from clinics differed in lowest illness-related BMI and age at lowest BMI from those in the community, in the US, using t-tests. We examined the construct validity for the online diagnostic questionnaire with the interview-based SCID by calculating the positive and negative predictive values for AN Criteria B and C, and for the presence of binge eating. Higher values correspond to increased accuracy of results. Furthermore, correlations for lowest illness-related BMI and age at low weight were calculated. Results are provided by country.

## Results

### General ANGI Descriptive Information

**Table 1** provides information regarding the number of cases and controls by site and source. Only those samples that passed quality control to be submitted for genotyping are included.

**Table 1.**
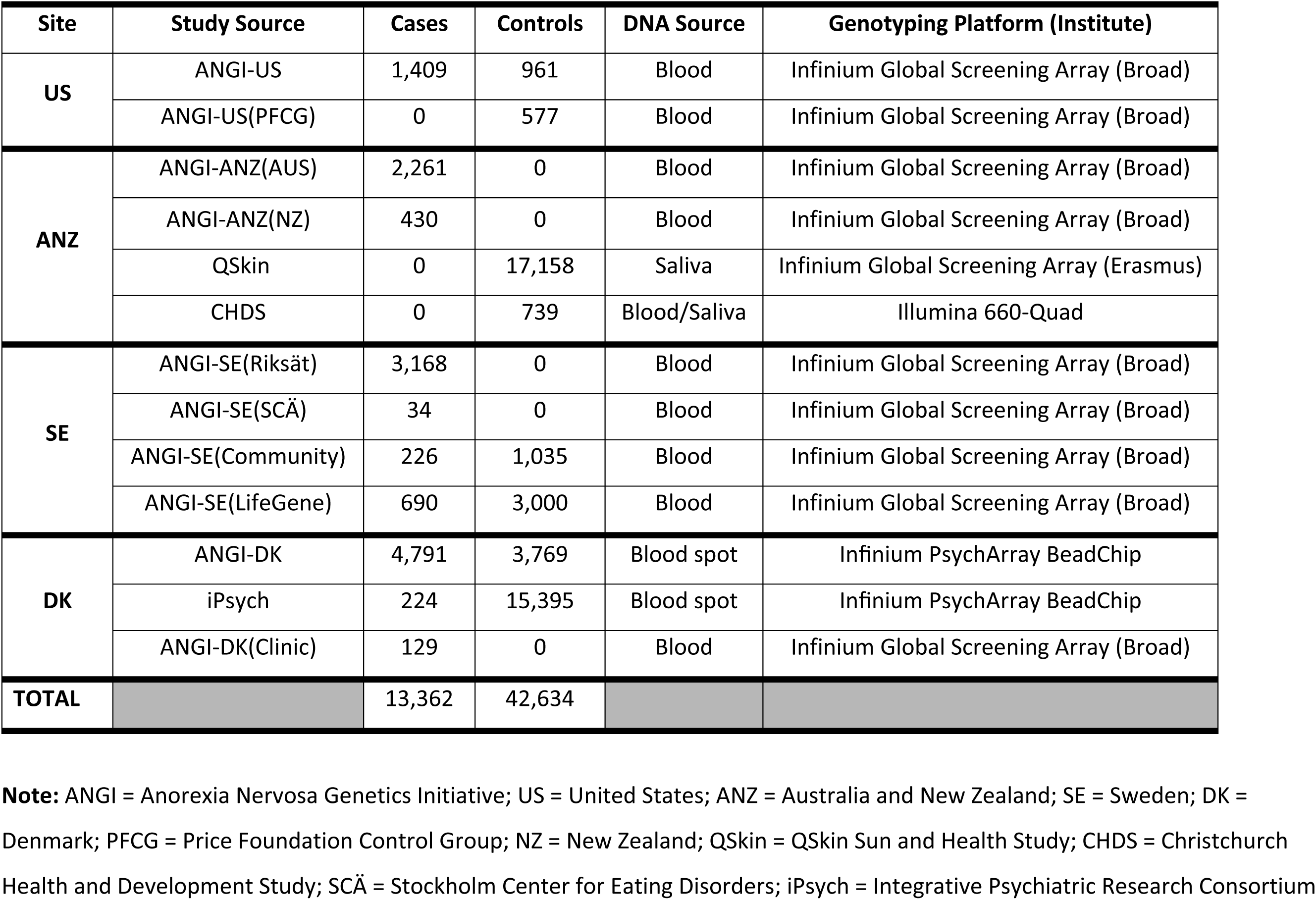
The number of samples collected and submitted for genotyping for ANGI, by site and source.

### Evaluation of Recruitment Sources

Details about where participants heard about ANGI in the US are provided in **Figure 2**. The most successful recruitment avenue for cases was the Internet, including social media, whereas, for controls, it was email followed by the Internet. Advocacy groups and clinicians (including clinical programs and eating disorder centers) were also important for recruiting cases. More than 20% of controls heard about ANGI through ResearchMatch (a not-for-profit online recruitment tool) or UNC (but not via email). Notably, 12% of cases and 13% of controls heard about ANGI from family members.

**Figure 2.**
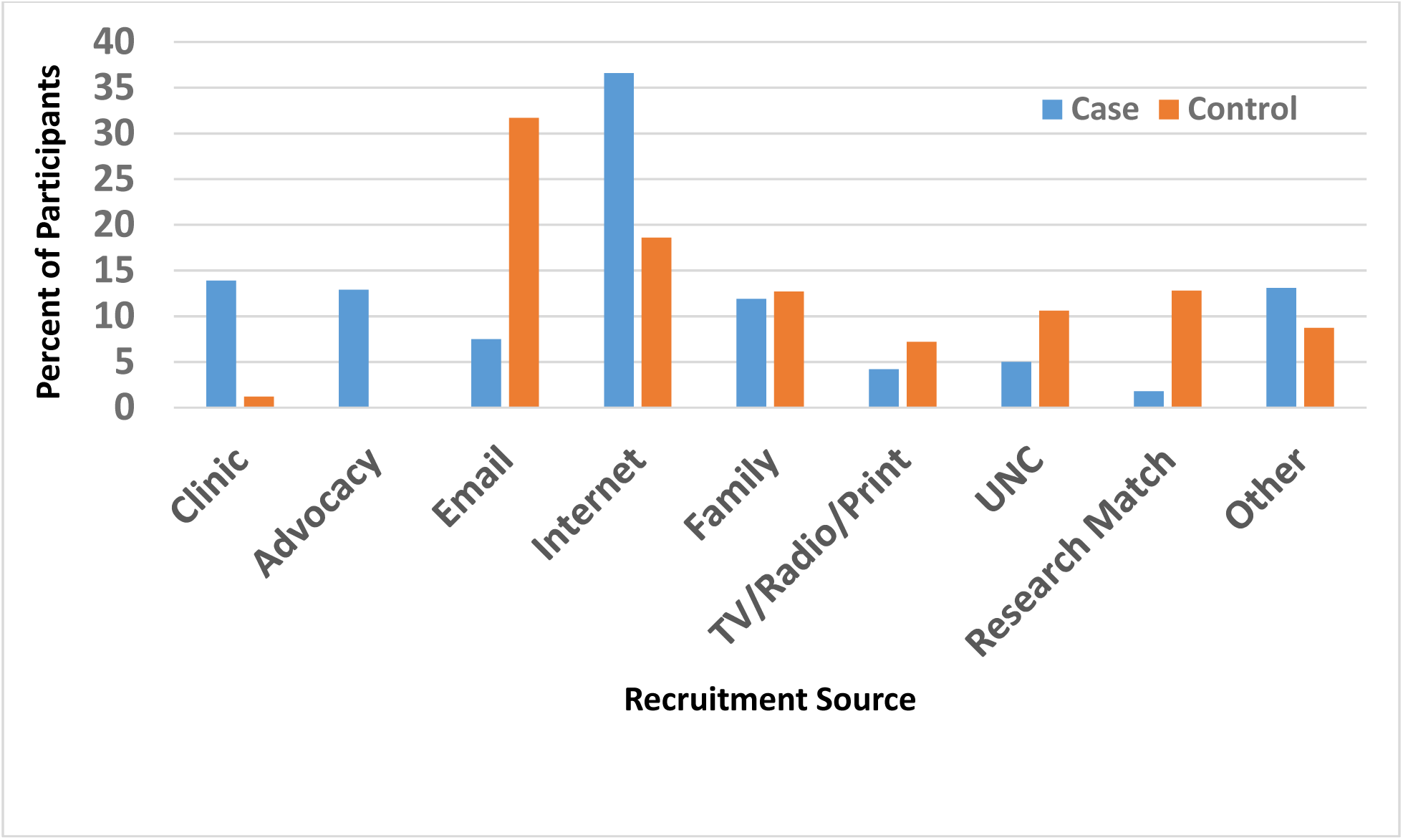
Sources of recruitment in the United States.?

**Figure 3** demonstrates the importance of using media outlets for participant recruitment. Although ANGI participation had increased steadily in Australia between the media launches in April 2013 and in March 2015, there was a tremendous spike in interest and study completion after the second launch. In approximately 2 months, the number of individuals who completed the ED100K-v1 questionnaire went from 2,228 to 3,574 individuals. Similarly, **Figure 4** illustrates the impact of media launches for ANGI-ANZ(NZ) recruitment.

**Figure 3.**
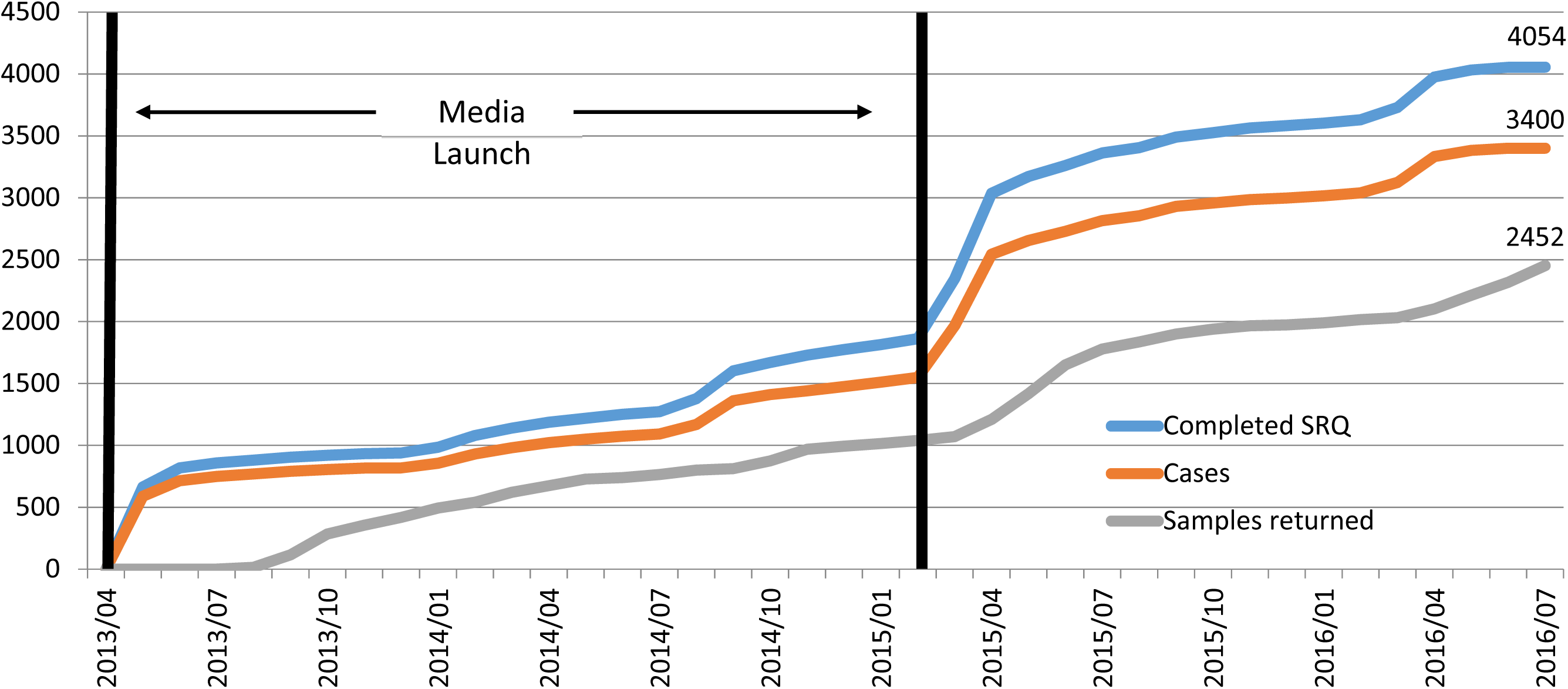
ANGI-ANZ(AUS) cumulative sample collection: progress after two media launches in Australia (April 2013 and March 2015).

**Figure 4.**
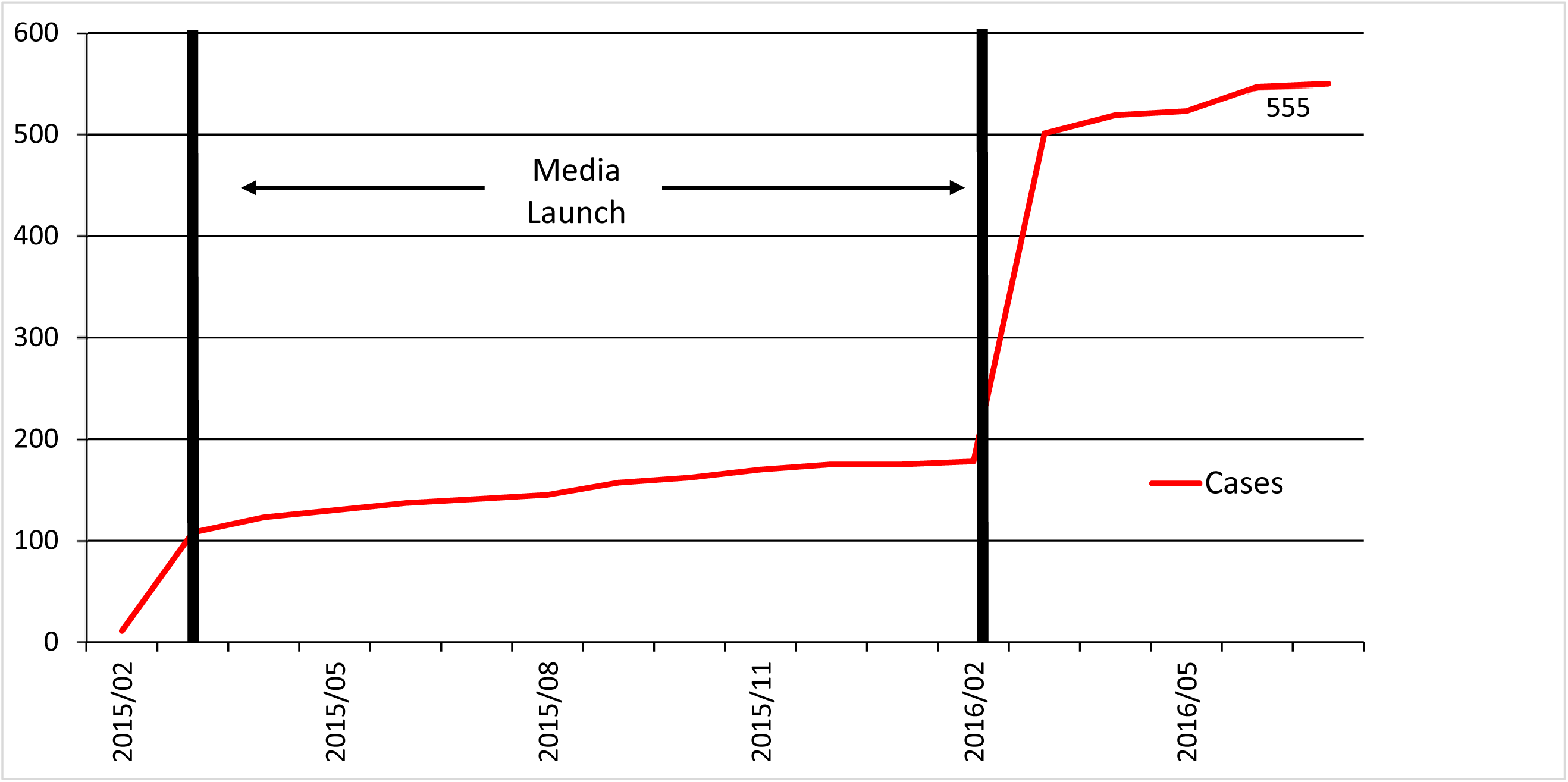
ANGI-ANZ(NZ) cumulative sample collection: progress after two media launches in New Zealand (March 2015 and March

We also explored whether cases from eating disorder treatment centers were significantly more ill than individuals from the community in the US. Cases ascertained through the community reported a significantly younger age at lowest weight [mean(sd)=17.4 (5.7)] than those recruited through eating disorders clinics [mean(sd)=19.5 (9.9)]. The two groups also differed significantly in lowest BMI with those from eating disorders clinics [mean(sd)=14.2 (2.1)], having lower BMIs than those from the community [mean(sd)=15.1 (1.9)]. However, the majority of participants in both groups, 82.8% from eating disorder clinics and 63.3% from the community, reported illness-related BMI values consistent with severe and extreme DSM-5 AN severity indices.

### Validity of Online Eating Disorder Questionnaire

**Table 2** lists the construct validity results. Positive predictive values were high in both countries, indicating that among those who had a positive screening test, the probability of AN Criterion B, Criterion C, and binge eating ranged from 85% to 100%. Results also indicated that among women who had a negative screening test, the probability of not having these AN criteria or binge eating was between 77% and 100%. For AUS, all participants endorsed Criterion C in the ED100K-v1 questionnaire and all but one individual endorsed Criterion C in the interview; thus, we could not calculate negative predictive value.

**Table 2.** Positive and negative predictive value (PV) of the ANGI online eating disorder questionnaire compared with the SCID-Module H (eating disorders).

|  | <u>United States (n = 51)</u> |  | <u>Australia (n = 58)</u> |  |
| --- | --- | --- | --- | --- |
|  | Positive PV | Negative PV | Positive PV | Negative PV |
| AN Criterion B | 1.00 | 1.00 | 0.96 | 1.00 |
| AN Criterion C | 0.98 | 1.00 | 0.98 | --- |
| Binge Eating | 0.85 | 0.77 | 0.93 | 0.83 |

Note: ANGI = Anorexia Nervosa Genetics Initiative; SCID = Structured Clinical Interview for the Diagnostic and Statistical Manual of Mental Disorders-IV; AN = anorexia nervosa. Criterion B is an intense fear of gaining weight or becoming fat, even though underweight. Criterion C is a marked disturbance in the way body weight or shape is experienced, undue influence of body weight or shape on self-evaluation, or denial of the seriousness of current low body weight.

The correlations between the ED100K-v1 questionnaire and interview responses for lowest illness-related BMI were high (US: *r*=0.91; AUS: *r*=0.92). The correlation for age at lowest weight in the US (*r*=0.58) was lower than that for AUS (*r*=0.90). In the US, the mean (SD) difference for age at lowest weight between the questionnaire and interview was 7.0 (10.7) years; three individuals reported a lower lifetime weight occurring between the self-report questionnaire and interview.

## Discussion

The primary goal of ANGI was to inform the etiology of AN by amassing the largest genetic sample of AN ever assembled and creating data and sample resources for future research. To accomplish this, we combined new and existing resources from the US, ANZ, SE, and DK. Although methods varied across countries, we were successful in ascertaining broadly representative cases and controls.

Our approaches optimized resources in each country. The existence of national patient registers (SE and DK) and quality registers (SE) facilitated identification of individuals who had been treated for AN (cases) as well as controls. The carefully curated biobank of PKU cards in DK made for a highly efficient ascertainment strategy, but precluded re-contacting participants for future research. The ability to link both the Swedish and Danish data to other national registers will allow us to expand analyses to include a broad range of phenotypes and exposures and even design gene-by-environment interaction investigations that include measured genotypes.

The majority of both clinic- and community-ascertained cases had illness-related BMIs in the DSM-5 severe and extreme ranges. However, those recruited via clinics reported older ages at low weight and somewhat lower illness-related BMIs than cases from the community. Notably, some cases from the community reported lowest BMI values that were lower than the range observed for those from the clinic. These observations are encouraging for general community recruitment for psychiatric genetics—at least for disorders in which there is strong social media presence. Although the percentage of individuals recruited via social media who had not received treatment is unknown, it raises ongoing concerns that many individuals with AN are distanced from clinical providers and not receiving care for their illness (13, 14, 31).

We were also able to provide information on the appropriateness of using a web-based diagnostic questionnaire for ascertainment of cases. ED100K-v1 operationalizes the DSM-IV criteria for all eating disorders (now adapted for DSM-5). The questionnaire was comparable to the SCID eating disorders module in determining the presence of criteria B and C for AN and in assessing lowest illness-related BMI and age at low weight in both the US and AUS. Given the need for large samples for successful GWAS, efficient case identification and minimally adequate phenotyping are methodologically important (7). A self-report questionnaire based on a structured interview that accurately identifies cases reduces the time and money spent on clinical interviewing and streamlines the case identification and ascertainment process.

ANGI does have limitations. First, ANGI participants are primarily of European ancestry. As yet, it is unclear whether psychiatric disorder GWAS findings can be generalized across populations (32). Second, AN primarily afflicts females. The small number of males recruited limits our capacity to determine sex differences in genetic effects. Future studies will require targeted strategies to ascertain and recruit substantial numbers of males. Third, ANGI used several ascertainment strategies to maximize recruitment in the shortest possible time. This could have led to the different sets of cases having different genetic architectures. However, the experience of the PGC (and other groups) provides strong evidence that, if this is true, the adverse impact is small relative to the large benefits of an increased sample (7, 33). Finally, the SE and DK collections enable linkage with other national registers that can provide a comprehensive characterization of psychiatric and somatic comorbidity, whereas these auxiliary data are not available in the US and ANZ.

In summary, the ability to generate large sample sizes for genomic investigations on eating disorders and other psychiatric disorders requires multi-site and international collaborations, as has been successfully demonstrated with the Wellcome Trust Case Control Consortium (34) and the PGC (6). The ANGI data are undergoing rigorous quality control and are being meta-analyzed with existing PGC-ED data. The combined sample size is approaching 20,0 cases with ancestrally matched controls, which should yield additional genome-wide signals.

Given the high prevalence of comorbid psychopathology in AN and eating disorders more broadly (13, 35, 36), these data can also be used for cross-disorder meta-analyses and SNP-based genetic correlations to identify putative genetic variants shared with other psychiatric and genetically influenced phenotypes. Moreover, an accurate and cost-efficient approach to case and control identification through self-report online questionnaires or existing national patient registers demonstrated here, is critical for rapid sample acquisition. Although ANGI was a highly successful collaboration, even larger sample sizes must be obtained to maximize GWAS findings. Ongoing collections from Europe and Asia are queued for genotyping, and large collections for other eating disorders (bulimia nervosa and binge-eating disorder) are underway. The PGC blueprint for the future aims for 100,000 cases for all major disorders is a scientifically justifiable aim (8). Uncovering the biological basis of eating disorders will aid in our understanding of their etiology, contribute to the development of novel or repurposed therapeutics, and ultimately reduce disability and mortality associated with the illnesses.

## Acknowledgements

The Anorexia Nervosa Genetics Initiative (ANGI) is an initiative of the Klarman Family Foundation. Additional support was from the NIMH Center for Collaborative Genomics Research on Mental Disorders, award U24 MH068457, and by the North Carolina Translational and Clinical Sciences Carolina Data Warehouse. Dr. Munn-Chernoff acknowledges funding from the National Institutes Health (NIH; K01 AA025113). Dr. Baker acknowledges funding from the National Institutes Health (NIH; K01 MH106675). Drs. Larsson, Petersen, and Mortensen acknowledges funding from an unrestricted grant from the Lundbeck Foundation, iPSYCH (The Lundbeck Initiative for Integrative Psychiatric Research), and by Aarhus University for CIRRAU (Centre of Integrated Register-Based Research). Dr. Watson acknowledges support from the National Institutes Health (NIH; U01 MH109528-02S1). Dr. Yilmaz acknowledges support from the National Institutes Health (NIH; K01 MH109782). Dr. Olsen acknowledge support from the National Health and Medical Research Council (NHMRC) Project Grant No. 1063061, Program Grant No. 1073898. Dr. Whiteman is supported by a Research Fellowship [APP1058522] from the National Health and Medical Research Council of Australia (NHMRC). Dr. Bergen is supported by a Professional Services Agreement with the Regents of the University of California. Dr. Kaplan received support from Ministry of Health of Ontario AFP Innovation Fund. Drs. Horwood and Boden acknowledge support from the Health Research Society of New Zealand (Programme Grant 16/600). Drs. Grove and Mattheisen acknowledge support from iPSYCH (The Lundbeck Foundation Initiative for Integrative Psychiatric Research). Drs. Jordan and Kennedy acknowledge funding from the University of Otago Research Grant. Dr. Wade acknowledges support from the National Health and Medical Research Council (NHMRC; 324715 and 480420). Dr. Landén acknowledges funding from the Swedish foundation for Strategic Research (KF10-0039). Dr. Sullivan acknowledges support from the National Institutes Health (NIH; R01 MH109528, D0886501) and the Swedish Research Council (Vetenskapsrådet, award D0886501). Dr. Bulik acknowledges funding from the Swedish Research Council (VR Dnr: 538-2013-8864).

The QSkin Study is supported by a Program Grant [APP1073898] and Project Grant [APP 1063061] from the National Health and Medical Research Council of Australia (NHMRC). The Australia & and New Zealand Academy for Eating Disorders also provided support for this investigation.

LifeGene was supported by the Ragnar and Torsten Söderberg Foundation, AFA Insurance, and the Stockholm County Council/Karolinska Institutet Research funds.

We acknowledge the support of the Price Foundation, Walden Behavioral Care, McCallum Place, and the Renfrew Center in assisting with recruitment. We express our gratitude to all of the bloggers who helped us disseminate information about ANGI, especially June Alexander http://www.junealexander.com/ and Carrie Arnold http://carriearnold.com/.

In Sweden, we acknowledge the assistance of the Stockholm Centre for Eating Disorders and wish to thank the Swedish National Quality Register for Eating Disorders: Riksät. We would also like to thank all the research nurses and data collectors at the Department of Medical Epidemiology and Biostatistics who have been working on ANGI.

In Australia and New Zealand, we would like to thank VIVA! Communications for their efforts in promoting the study, and the Butterfly Foundation and June Alexander for their ongoing support of anorexia nervosa research in Australia and EDANZ in New Zealand. We also acknowledge the assistance of Dr. Sarah Maguire and Professor Janice Russell (University of Sydney), Professor Phillipa Hay (Western Sydney University), Dr. Sloane Madden (Western Sydney University and the Sydney Children’s Hospital Network), Professor Susan Sawyer and Dr. Elizabeth Hughes (Royal Children’s Hospital, Melbourne), Dr. Kate Fairweather-Schmidt (Flinders University), Dr. Anthea Fursland (Centre for Clinical Interventions and Curtin University), Julie McCormack (Princess Margaret Hospital for Children), Dr. Fiona Wagg (Royal Hobart Hospital), and Dr. Warren Ward (Royal Brisbane and Women’s Hospital) in recruitment. We also thank Lorelle Nunn for her validation work on the ED100Kv1 Questionnaire. Additionally, administrative support for data collection was received from the Australian Twin Registry, which is supported by an Enabling Grant (ID 310667) from the NHMRC administered by the University of Melbourne. In New Zealand, we also acknowledge assistance with recruitment from Dr. Marion Roberts (University of Auckland), Rachel Lawson (South Island Eating Disorders Service), Michelle Meiklejohn (Auckland District Health Board), and Dr. Roger Mysliwiec. Special thinks to those who provided their stories in relation to publicity about ANGI.

We are deeply grateful to all of the individuals who, through their participation, made ANGI a success. The goodwill that permeated the eating disorders community fueled by the enthusiasm of prominent bloggers, advocates, clinicians, treatment centers, scientists, organizations, families, and especially those who have suffered from anorexia nervosa, yielded in an unprecedented and inspired global movement to complete this science.

## Footnotes

1 In Australia, the interviewer reviewed the SCID protocol and instructions, observed SCID interviews, and reviewed existing interviews with staff from the US. Although the interviewer was not observed administering a SCID by a PhD-level clinician in the United States, this individual did have nearly 15 years of experience administering diagnostic interviews, including the SCID.

